# Identification and characterization of metabolic subtypes of endometrial cancer using systems-level approach

**DOI:** 10.1101/2023.01.05.522818

**Authors:** Akansha Srivastava, P K Vinod

## Abstract

**Background:** *Endometrial cancer* (EC) is the most common gynaecological cancer worldwide. Understanding the metabolic adaptation and its heterogeneity in tumor tissues may provide new insights and help in cancer diagnosis, prognosis, and treatment. In this study, we investigated metabolic alterations of EC to understand the variations in the metabolism within tumor samples.

**Methods:** We integrated the TCGA transcriptomics data of EC (RNA-Seq) with the human genome-scale metabolic model (HMR2.0) and performed unsupervised learning to identify the metabolic subtypes of EC and uncover the underlying dysregulated metabolic pathways and reporter metabolites in each subtype. The relationship between metabolic subtypes and clinical variables was explored. Further, we characterized each subtype at the molecular level and correlated the subtype-specific metabolic changes occurring at the transcriptome level with the genomic alterations.

**Results:** EC patients are stratified into two robust metabolic subtypes (cluster-1 and cluster-2) that significantly correlate to patient survival, tumor stages, mutation, and copy number variations. We observed coactivation of pentose phosphate pathway and one-carbon metabolism along with genes involved in controlling estrogen levels in cluster-2, which is linked to poor survival. PNMT and ERBB2 are also upregulated in cluster-2 samples and present in the same chromosome locus 17q12, which is amplified. PTEN and TP53 mutations show mutually exclusive behavior between subtypes and display a difference in survival.

**Conclusion:** This work identifies metabolic subtypes with distinct characteristics at the transcriptome and genome levels, highlighting the metabolic heterogeneity within EC.

## Background

Metabolic reprogramming is one of the emerging hallmarks of cancer [1]. Different studies have shown that altered metabolism relates to cancer development, progression, and outcomes [2]. Cancer cells rewire their metabolism to support their rapid growth and division. Metabolic pathways are dysregulated to fulfil the increased demand for nutrients and energy. An adequate balance between nutrient metabolism and energy production helps cancer cells to maintain homeostasis. The cancer cell metabolism is affected by different intrinsic factors, such as epigenetics, gene expression, cellular composition, altered signaling pathways, and microbial populations [3]. Metabolic reprogramming in cancers is driven by activated oncogenes and/or inactivated tumor suppressor genes [4]. It is also influenced by extrinsic factors like tumor microenvironment, obesity, diet, and genetic factors [3]. Metabolic phenotypes transform over time as tumor growth progresses from early-stage to late-stage [3]. All these factors are responsible for metabolic heterogeneity, resulting in the diverse metabolic profiles of cancer patients. Therefore, tumor tissues adopt different mechanisms to reprogram the metabolic pathways, depending on tissue types, mutation patterns (gene amplification or deletions), and other cellular preferences. Some cancer types favor glycolysis over oxidative phosphorylation for energy production, known as the Warburg effect [5]. Comprehending the various aberrant metabolic processes and their fundamental causes can provide new insights and helps in the diagnosis, prognosis, and treatment of cancer.

*Endometrial cancer* (EC) is the most common gynaecological cancer worldwide, which emerges in the inner lining of the uterus and may spread to the outside of the uterus in advanced stages [6]. The number of EC cases increases by 1% annually in the United States [7]. The death rate has increased by 2% per year over the last decade, and there is no improvement in the survival rate of EC. The possible reasons could be decrease in fertility rate, obesity, and sedentary lifestyles [7]. Based on clinical and epidemiological studies, EC is generally divided into Type 1 and Type 2 endometrial carcinomas [8]. The endometrioid endometrial Adenocarcinomas and serous endometrial adenocarcinomas are the two most common histological subtypes of EC. Type 1 EC comprises endometrioid subtypes. It is less aggressive, characterized by high estrogen levels, and has better survival outcomes [9]. In contrast, Type 2 EC includes serous carcinoma. It is more aggressive, independent of estrogen levels, and has a poor survival rate [9]. Levine et al., (2013) comprehensively analyzed the genomic, transcriptomic, and proteomic profiles of 363 EC patients. They identified four molecular subtypes of EC: POLE ultramutated, microsatellite instability hypermutated, copy-number low, and copy-number high [10]. Recent studies indicate that the heterogeneous metabolic profile of cancer patients influences their survival outcomes [11]. Therefore, it is crucial to stratify the EC patients based on their metabolic profile, which may help discover the underlying aberrant metabolic processes and design therapeutics for treatment.

Transcriptomic analysis of metabolic pathways has been carried out to elucidate the reprogramming of metabolism in different tumor tissues. Peng et al., (2018) analyzed the mRNA expression of seven metabolic pathways in 33 cancer types. They revealed that upregulation of carbohydrate, nucleotide, and vitamin/cofactor metabolism is consistently related to a worse prognosis, while upregulated TCA cycle and lipid metabolism correlates with a better prognosis [12]. Another pan-cancer study by Rosario et al., (2018) examined the metabolic expression profile of the tumor-matched normal and tumor samples across 26 cancer tissues and uncovered common and unique metabolic characteristics of major cancer types [13]. In addition, Sheraj et al., (2021) adopted a more targeted approach to explore the metabolic changes using the transcriptomic data of 20 tumor types and revealed that genes involved in central metabolism have significant transcriptional alterations [14]. Zhang et al., (2020) performed a co-expression analysis of metabolic pathways in 33 cancer types and identified folate metabolism, glutamine metabolism, glycine and serine metabolism, and purine nucleotide metabolism to be altered between normal and tumor samples [15]. These pan-cancer studies focused on discovering metabolic rewiring across cancer types. Trousil et al., (2014) demonstrated that choline metabolism is altered in EC using the nuclear magnetic resonance (NMR) technique [16]. Byrne et al., (2014) have compared the normal endometrium with tumor samples and observed upregulation of glycolytic–lipogenic metabolism in EC [17]. Glutathione metabolism is also dysregulated in EC [18].

These studies are limited to metabolic alterations between normal and tumor samples. The metabolic heterogeneity among EC patients remains unclear and has not been studied yet.

In this work, we combined the transcriptomic profile of patients with the human genome-scale metabolic model (HMR2.0) to identify the metabolic subtypes of EC. We also examined the characteristics of each subtype at the genomic level. Our study provides subtype-specific biomarkers and reporter metabolites, which will be helpful in the diagnosis and prognosis of EC.

## Methods

### Datasets and Data Pre-processing

We downloaded the TCGA RNA-Seq data (HTSeq Count) of EC using TCGAbiolink [19]. The TCGA MC3 files were retrieved from the Genomic Data Commons (GDC) portal to analyze the mutation profile of EC patients. The cBioportal was used to obtain the segmented copy number variation datasets [20]. We considered TCGA-Clinical Data Resource (CDR) to retrieve the corresponding clinical annotation for EC patients [21]. For validation, the microarray series dataset (GSE17025, Affymetrix Human Genome U133 Plus 2.0 Array), consisting of 91 EC samples, was downloaded from the NCBI GEO database using the GEOquery package [22]. We employed the human genome-scale metabolic model (HMR2.0) to study cancer metabolism. The model consists of 8181 reactions, 6006 metabolites, and 3765 genes, which describe the standard metabolic processes of a human cell.

The TCGA RNA-Seq dataset consists of 565 samples, out of which 542 are primary tumor samples (sample type code - 01) and 23 are tumor-matched-normal samples (sample type code - 11). We used the variance-stabilizing transformation (VST) to normalize RNA-Seq raw count data. The GEO microarray dataset consists of 79 endometrioid and 12 papillary serous samples with various grades. We used the microarray data normalized using Affymetrix’s MAS5.0 algorithm and log_2_ transformed.

### Identification of Clusters

To identify the metabolic subtypes, the transcriptomic data was combined with the Human genomescale metabolic model. Out of 3765 metabolic genes, 3584 were present in the gene expression dataset. The median absolute deviation (MAD) was computed for all metabolic genes, and the top 1000 genes based on the MAD score were selected for clustering. We adopted a consensus clustering approach to identify robust clusters. The non-negative matrix factorization (NMF) method was applied to cluster the 542 EC patients using the NMF R package [23]. The silhouette width, dispersion, and cophenetic correlation coefficient were utilized to decide the optimal number of clusters. We also experimented with six NMF methods (brunet, KL, lee, Frobenius, offset, nsNMF). The performance of all methods was evaluated on the same criteria used for determining the number of clusters.

### Differential Gene Expression Analysis

The differential gene expression analysis was performed using the DESeq2 to identify the different metabolic alterations between clusters [24]. To study the metabolic changes that occur from normal to tumor conditions, we performed differential gene expression analysis between 23 pairs of tumor-matched normal and tumor samples. The differentially expressed genes (DEGs) were identified based on log_2_ fold change (FC) (|log_2_FC|) > 1 and adjusted p-value (adj p-value) < 0.05. For the GEO dataset, we identified DEGs using limma R package with criteria of |log_2_FC| > 0.6 and adj p-value < 0.05 [25].

### Genomic Profile of Clusters

The TCGA somatic mutation data and copy number variation (CNV) datasets were used to assess the unique characteristics of clusters at the genome level. The number of somatic mutation and CNV samples was 525 and 534, respectively. The “maftool” was employed to analyze and visualize the mutation profile of clusters [26]. The differentially mutated genes between clusters were identified based on the minimum mutation frequency of 10 using the Fisher exact test (p-value < 0.01). We used the Genomic Identification of Significant Targets in Cancer (GISTIC) 2.0 method to identify the significant recurrent CNVs specific to each cluster [27]. The analysis was performed with amplification/deletion threshold > |0.1|, a confidence interval of 0.99 and q-value < 0.05. The GISTIC results were also visualized with maftool.

### Reporter Metabolite Analysis

The mapping of significant transcriptional changes in the metabolic network facilitates the identification of highly regulated metabolites, known as reporter metabolites. We performed reporter metabolite analysis by integrating the p-value and log_2_FC score obtained from DESeq2 analysis with the human genome-scale metabolic model. The reporter metabolite algorithm first utilizes an inverse normal cumulative distribution to convert the p-value of an enzyme into the z-score [28]. Then, the method identifies the enzymes around each metabolite and computes the z-score of that metabolite using the following equation:

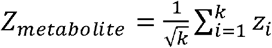

Here, k represents the number of enzymes around each metabolite. 100000 sets of k enzymes were chosen randomly to correct the Z-score of each metabolite by subtracting the mean (*μ_k_*) and dividing by the standard deviation (*σ_k_*) of the aggregated Z scores of all sets. The corrected Z-score was transformed into a p-value. We selected reporter metabolites with a minimum of 3 neighboring genes and a p-value < 0.05.

### Association of Clusters to Clinical Information

The log-rank test was performed to assess the survival differences between clusters. The Kaplan Meier Plot was generated using R package. The relationship of clusters with the clinical variables such as clinical stage, histological type, histological grade, and age was determined by the Fisher exact test. Cramer’s V was also computed to quantify the association of the clinical variables with the clusters. Its values vary from −1 to +1. A value of +1 dictates the stronger association; 0 represents no association, and −1 represents a strong association in the opposite direction. In addition, the univariate cox regression analysis was performed to determine the DEGs significantly associated with the survival outcomes of EC patients. A p-value < 0.05 was considered for identifying prognostic genes.

## Results

To study metabolic reprogramming in EC, we integrated the gene expression data of EC with the human genome-scale metabolic model, which consists of 3765 metabolic genes. First, we performed the principal component analysis (PCA) to comprehend the metabolic profile of EC patients. PCA plots revealed that normal samples have different metabolic gene expression profiles compared to tumor samples (**Figure 1a**). It also indicates that the metabolic profile of EC patients is heterogeneous.

**Figure 1:**
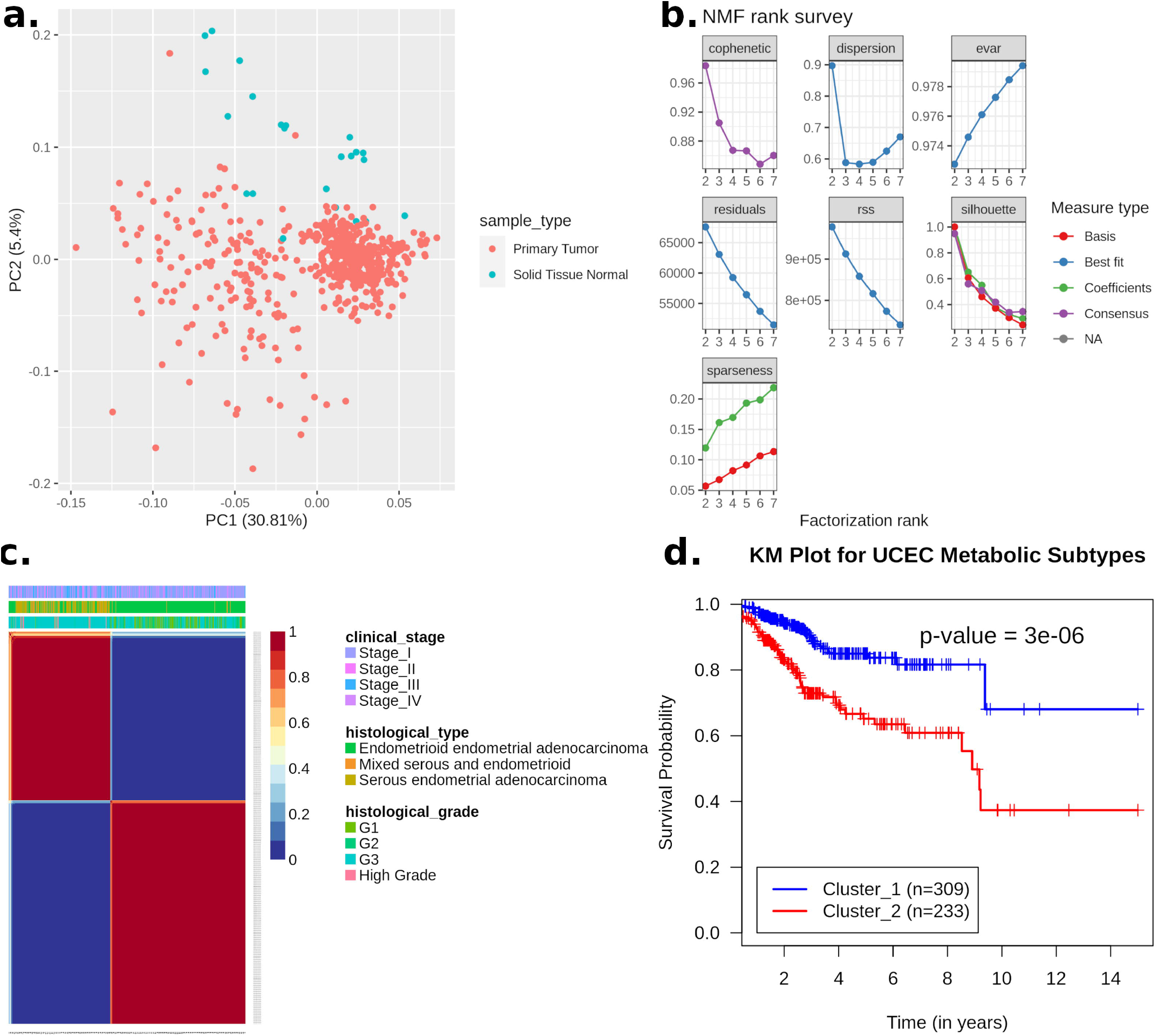
Clustering of EC patients based on the metabolic gene expression profile. (a) Principal component analysis reveals metabolic heterogeneity within EC. The x-axis represents the first principal component, which captures 30.81% variation, and the y-axis represents second component, which captures 5.4% variation. (b) Identifying an optimal number of clusters from different cluster sizes (k = 2-7). The y-axis represents the evaluation metric, and the x-axis denotes the number of clusters. (c) Consensus plot showing two robust clusters cluster-1 and cluster-2 (silhouette width = 0.98, cophenetic coefficient = 0.996 and dispersion = 0.96) of EC patients. The plot also includes information on different clinical variables. (d) Kaplan Meier Plot shows metabolic subtypes have significantly different survival probability (p-value < 0.001).

To discover the metabolic subtypes within EC, we applied the NMF method to cluster the EC patients using their gene expression profile of 1000 metabolic genes filtered based on MAD score. Different cluster sizes (2 to 7) were experimented with the brunet algorithm to tune the optimal number of clusters (**Figure 1b**). We identified two as the optimal number of the cluster based on silhouette width, dispersion, and cophenetic correlation coefficient. The Offset NMF technique among the different variants of NMF methods achieved the best performance (silhouette width = 0.98, dispersion= 0.96, cophenetic= 0.995) to classify the 542 EC patients into two subtypes: cluster-1 consists of 309 patients and cluster-2 has 233 patients. We ran the offset algorithm with default parameters for up to 2000 iterations and repeated the experiment 50 times, which yielded two robust clusters (**Figure 1c**). To assess the survival difference between clusters, we performed an overall survival analysis. Both clusters showed significant differences in survival outcomes (p-value < 0.001). Cluster-1 is associated with better survival, while cluster-2 is associated with poor survival (**Figure 1d**).

**Figure 2** shows the cluster-wise distribution across stages, histological types, grades, and age. The Fisher exact test shows that both clusters are significantly related to the clinical variables (p-value < 0.05) (**Table S1**). Cluster-1 is dominated by stage 1 samples, whereas Cluster-2 is dominated by stage 3 and stage 4 samples (**Figure 2a**). Stage 2 samples are almost equally distributed into both clusters. The Cramer’s V value of 0.36 shows a moderate association between clinical stage and clusters. Considering histological types, cluster-1 mainly consists of endometrioid endometrial adenocarcinoma samples, while cluster-2 comprises all histological types (**Figure 2b**). Both serous and endometrioid types have an almost equal distribution in cluster-2 samples. The Cramer’s V value of 0.61 describes strong relationships between clusters and histological types. In terms of tumor grade, cluster-1 consists primarily of grade 1 to grade 3 samples (**Figure 2c**). However, the distribution of grade 3 samples is more in cluster-2 compared to cluster-1. All high grades samples only belong to cluster-2. The Cramer’s V value for a histological grade is 0.58, which indicates a strong relationship between them. In addition, we divided the samples into two categories to perform the Fisher exact test between age and clusters (< 50 years and ≥ 50 years). The association of age with clusters is significant (p-value = 0.02), but Cramer’s V value for age is very small (0.098), indicating a negligible association of clusters with age (**Figure 2d**).

**Figure 2:**
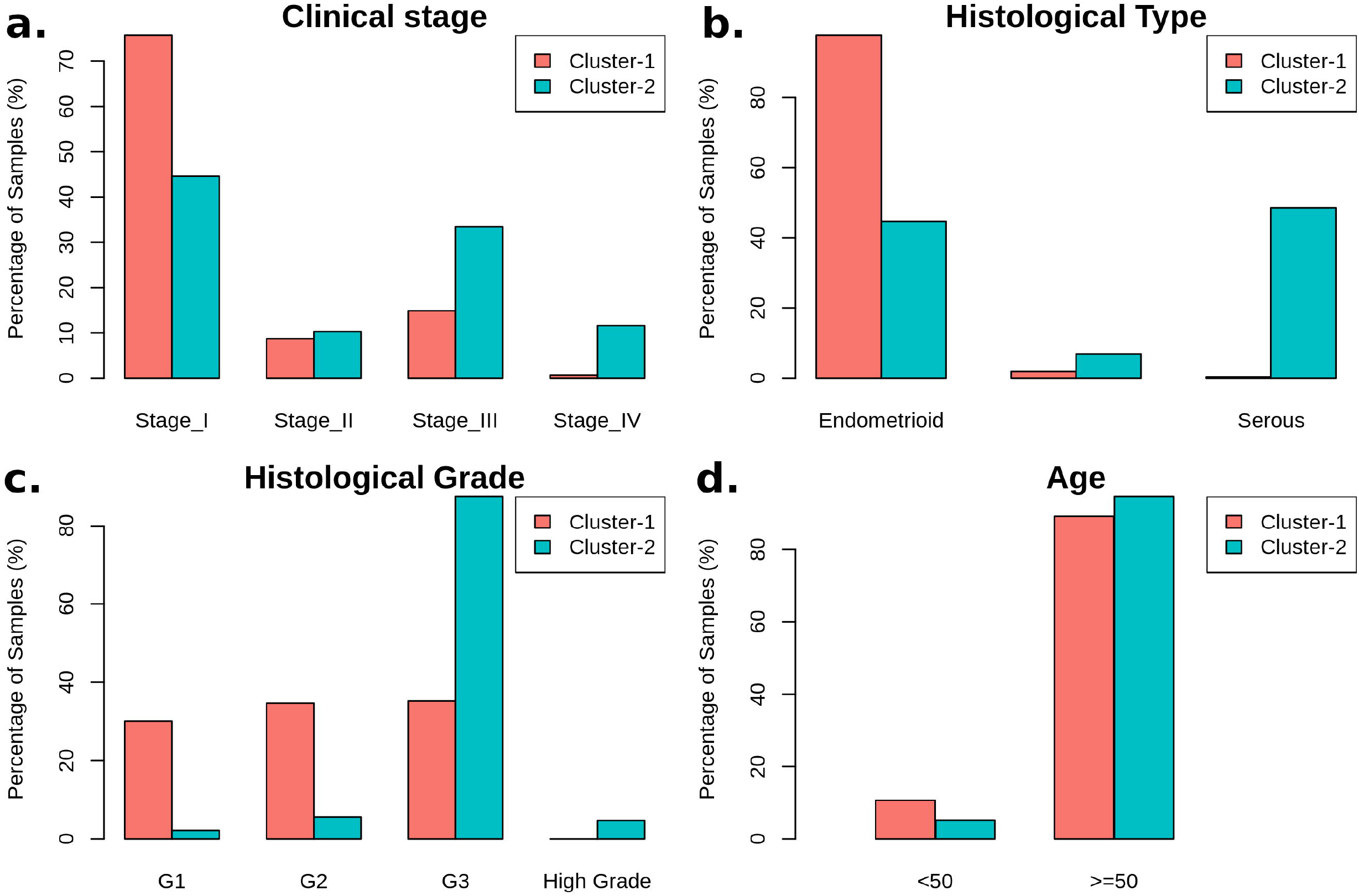
The bar plot showing the distribution of clinical variables in each cluster. (a) Clinical Stage, (b) Histological Type, (c) Histological Grade and (d) Age.

### Metabolic gene alterations in EC

To identify the unique metabolic characteristics of clusters at the transcriptome level, we performed differential gene expression analysis between cluster-1 and cluster-2, considering the expression of 3584 metabolic genes. There are 264 upregulated and 137 downregulated genes between cluster-1 and cluster-2 samples based on criteria of |log_2_FC| > 1 and adj p-value < 0.05 (**Data S1**). The top candidate genes are shown in **Figure 3a**. We also conducted differential gene expression analysis between matched normal and tumor samples to uncover the common metabolic transcriptional changes in EC. We identified 971 DEGs (517 upregulated and 454 downregulated) between normal and tumor samples (**Data S1**). We compared the DEGs of both analyses and found that there are only 46 genes that are upregulated in both cluster-2 and tumor samples (**Figure 3b**). Most of the genes (151) are uniquely upregulated in cluster-2, which indicates a different set of genes responsible for cancer progression in cluster-2 samples.

**Figure 3:**
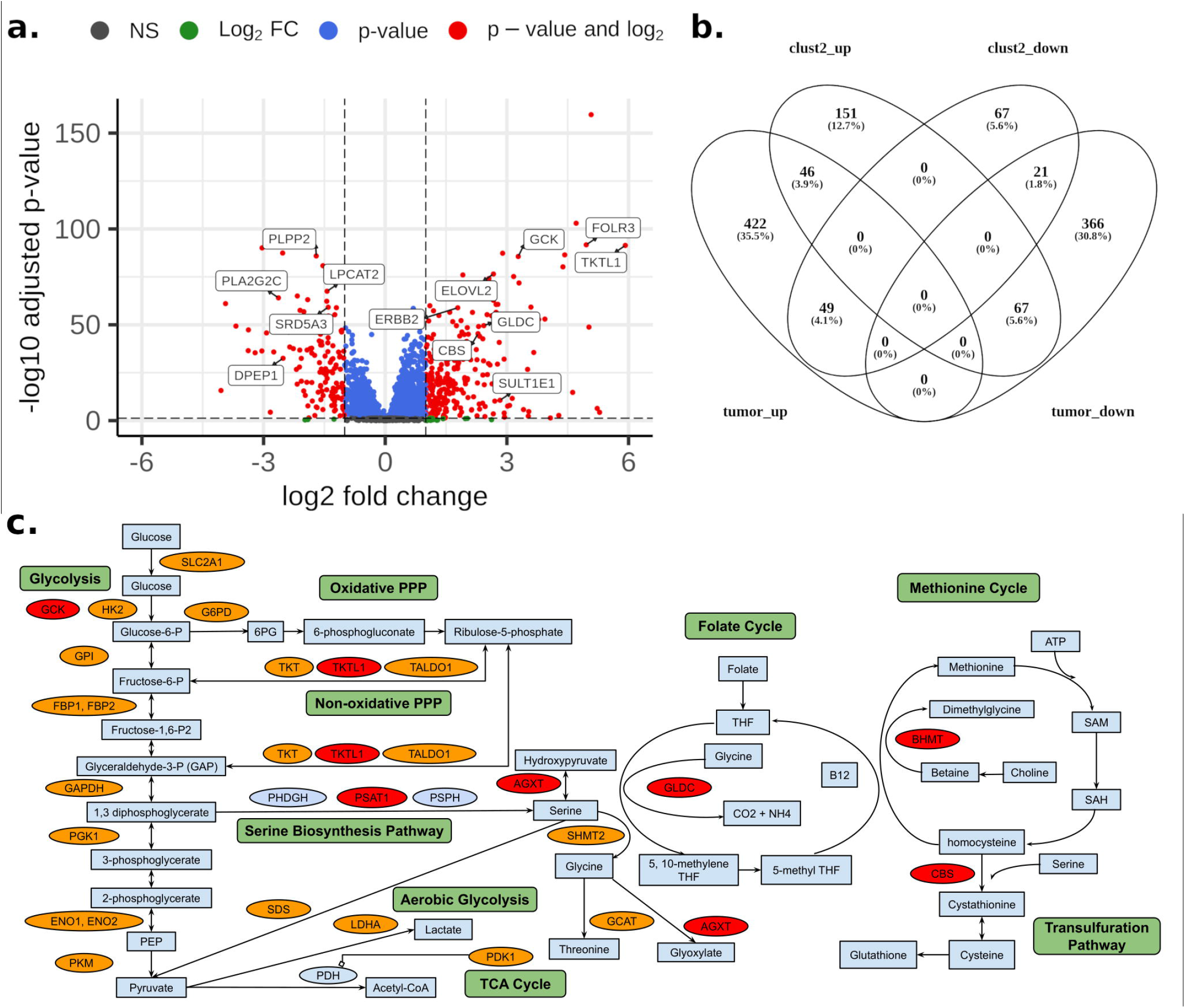
(a) Volcano plot showing DEGs between cluster-1 and cluster-2 samples. (b) Venn diagram showing the overlap of DEGs between cluster-1 vs cluster-2 and normal vs tumor comparisons (clust2_up: upregulated genes in cluster-2, clust2_down: downregulated genes in cluster-2, tumor_up: upregulated genes in tumor, tumor_down: downregulated genes in tumor). Venny 2.1 was used to generate the venn diagrams. (c) Metabolic pathways altered in EC and cluster-2 samples. Orange and red colors represent upregulated genes in normal vs. tumor conditions and cluster-1 vs. cluster-2 conditions, respectively.

Genes of the glycolysis pathway (SLC2A1, HK2, GPI, FBP1, FBP2, PFKFB1, PFKFB2, PFKFB4, TPI1, GAPDH, PGK1, ENO1, ENO2, PKM) are upregulated in EC (**Figure 3c**). PDK1 inhibits oxidative decarboxylation of pyruvate by PDH and is upregulated in EC samples along with LDHA, which controls the interconversion of pyruvate and lactate. Further, two critical enzymes of the PPP, G6PD, which catalyzes the conversion of glucose-6-phosphate into 6-phosphogluconolactone, and PGD, which yield ribulose 5-phosphate by catalyzing the oxidative decarboxylation of 6-phosphogluconate, are upregulated in EC samples. PPP plays a role in the synthesis of nucleotides and generating coenzymes to maintain redox homeostasis. Further, enzymes of the non-oxidative phase of the PPP (TKT, TALDO1) are also upregulated in EC samples. Interestingly, TKTL1, another critical enzyme that connects the PPP with glycolysis by converting D-xylulose 5-phosphate into D-glyceraldehyde 3-phosphate, is upregulated in cluster-2 samples (log_2_FC = 5.91).

We observed that genes (BHMT, PSAT1, AGXT, DAO, GLDC, CBS) involved in glycine, serine, and threonine metabolism are upregulated in cluster-2 samples (**Figure 3c**). PSAT1 catalyzes the conversion of 3-phosphohydroxypyruvate to phosphoserine, which is further used in the formation of serine. Downregulation of PSAT1 promotes growth inhibition and induces apoptosis [29]. Cells with high PSAT1 levels increase the ratio of GSH (reduced) to GSSG (oxidized) and lower ROS. GLDC catalyzes the conversion of glycine to 5,10-methyleneteTHF, which is an essential intermediate in the folate cycle. CBS, a vital gene of the transsulfuration pathway, catalyzes the conversion of serine to cystathionine, the precursor of L-cysteine. BHMT catalyzes the conversion of homocysteine to methionine and dimethylglycine using choline-derived betaine as a methyl donor. We also observed genes of glycine, serine, and threonine metabolism (GLDC, PSAT1, SHMT2, GCAT, and SDS) to be upregulated in EC samples. However, the expression of GLDC and PSAT1 is very high in cluster-2 samples.

Estrogen sulfotransferase enzyme SULT1E1 is upregulated in cluster-2 samples. SULT1E1 catalyzes the sulfation of estrogens (estradiol and estrone) and catechol estrogens. UGT1A1 of the glucuronidation pathway is upregulated in cluster-2 samples. It encodes the UDP-glucuronosyltransferase enzyme, a major contributor to the glucuronidation activity associated with estrogens and catechol estrogens. Cytochrome P450 gene CYP1A1 is also upregulated in cluster-2 samples. CYP1A1 catalyzes the 2-OH hydroxylation of estrogens. This suggests that cluster-2 may be less related to estrogen exposure compared to cluster-1 samples. 17β-hydroxysteroid dehydrogenases HSD17B7 and HSD17B2, involved in the interconversion between estrone and estradiol, are upregulated in EC samples. On the other hand, CYP1B1, which catalyzes the 4-OH hydroxylation of estrogens, is downregulated in EC samples. The glutathione S-transferase enzymes (GSTA1, GSTA2, GSTA3, GSTM1, GSTM2, GSTM3, GSTM5) are also downregulated in EC samples. GSTs play a role in the inactivation of catechol estrogen. These observations indicate that estrogen exposure plays a critical role in the development of EC.

On the other hand, we observed genes (GGT1, GGCT, OPLAH, ANPEP, G6PD, PGD, IDH1, IDH2, GPX2, GPX7, GSR) that control the intracellular content of glutathione through *de novo* synthesis and regeneration of glutathione (GSH) from Glutathione disulfide (GSSG) to be upregulated in EC samples. GGT1, GGCT, OPLAH, and ANPEP play critical roles in synthesizing glutathione from glutamate, cysteine, and glycine. Similarly, the upregulation of G6PD, PGD, IDH1, and IDH2 can increase NADPH which is required to reduce glutathione. GPX2 and GPX7 are involved in converting GSH to GSSG to reduce the hydrogen peroxide and superoxide radicals, while GSR catalyzes the reduction of GSSG to GSH.

The expression of genes in the urea cycle (ASS1, CPS1, OTC, SLC25A13, and SLC25A15) is altered in EC samples. Urea cycle dysregulation promotes nitrogen utilization for pyrimidine synthesis [30]. Downregulation of ASS1 expression promotes cancer proliferation by diversion of its aspartate substrate toward the pyrimidine synthesis pathway [31,32]. SLC25A13, which exports aspartate from the mitochondria, is upregulated in EC samples.

### Reporter Metabolites

Further, we performed reporter metabolite analysis to identify altered metabolites between cluster-1 and clusters-2 and normal and tumor samples. The analysis was done with all DEGs, upregulated DEGs, and downregulated DEGs in cluster-2 and all tumor samples. We mapped the reporter metabolites with HMR2.0 metabolic pathways to infer the altered pathway. The metabolites of one-carbon metabolism (betaine, Choline, Glycine, serine, threonine, homocysteine, methionine, and THF) and the PPP (fructose-6-phosphate, ribose-5-phosphate, D-Xylulose-5-phosphate, and erythrose-4-phosphate) are reporter metabolites of cluster-2 samples (**Figure 4**). Both one-carbon metabolism and PPP are at the crossroads of anabolism and redox homeostasis. We also found glutamine as a reporter metabolite in cluster-2 samples. In addition, the metabolites of fatty acid synthesis, fatty acid elongation, omega-3 fatty acid metabolism, and omega-6 fatty acid metabolism are associated with upregulated genes of cluster-2 samples (**Figure S1**). In EC samples, we observed SAM and SAH as reporter metabolites along with metabolites of glycolysis (fructose-6-phosphate, fructose-2,6-bisphosphate, Glyceraldehyde-3-phosphate (GAP)), PPP (ribose-5-phosphate, ribulose-5-phosphate), TCA cycle (succinyl-CoA, isocitrate, oxalate) and lysine metabolism (L-lysine, N6,N6-dimethyl-L-lysine, N6,N6,N6-trimethyl-L-lysine, Histone-L-lysine, histone-N6-methyl-L-lysine). On the other hand, the metabolites associated with downregulated genes of cluster-2 samples map to phenylalanine, tyrosine, and tryptophan biosynthesis, sphingolipid metabolism, and glycerophospholipid metabolism (**Figure 4** and **Figure S1**).

**Figure 4:**
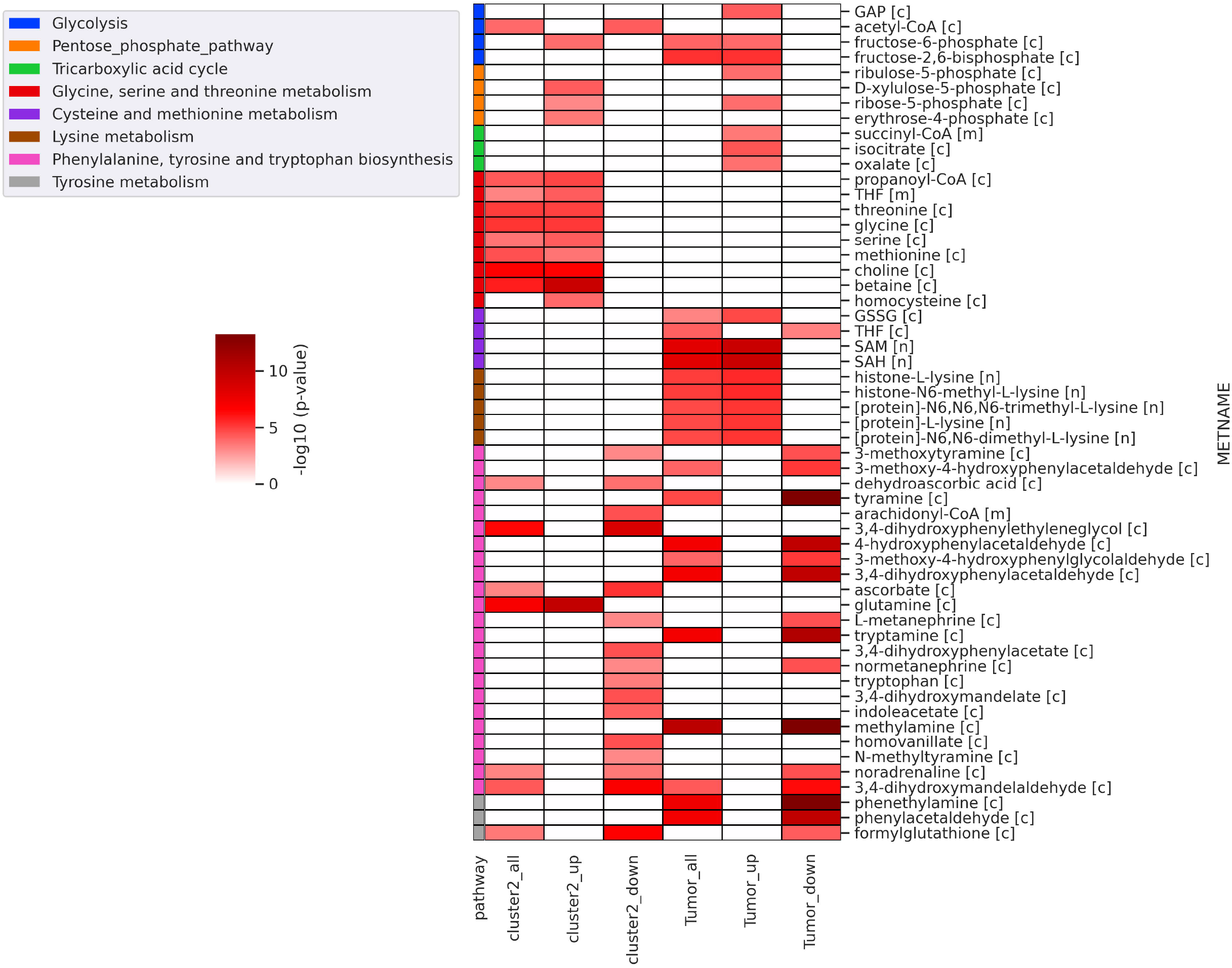
Heatmap of significant reporter metabolites involved in various metabolic pathways obtained for different conditions (cluster2_all: all DEGs of cluster-1 vs. cluster-2 condition, cluster2_up: upregulated genes in cluster-1 vs. cluster-2 condition, cluster2_down: downregulated genes in cluster-1 vs. cluster-2 condition, tumor_all - all DEGs of normal vs. tumor conditions, tumor_up: upregulated genes in normal vs. tumor conditions, tumor_down: downregulated genes in normal vs. tumor conditions). The metabolite name (METNAME) also includes the compartment information, [c] - cytosol, [m] - mitochondria, [n] - nucleus.

### Prognostic metabolic genes of EC

We performed a univariate cox regression analysis to discover the prognostic genes, which are differentially expressed between clusters. We observed that 225 DEGs in cluster-2 samples are associated with the survival of EC patients (**Data S1**). Among them, the high expression of 156 genes is associated with poor survival (Hazard ratio > 1), while the high expression of 69 genes is related to better survival outcomes (Hazard ratio <1). There are 86 genes that are only upregulated in cluster-2 samples and have a significant association with survival outcomes. These include genes of amino acid metabolism (PNMT, DDC, BHMT, CBS, TH, GLS), estrogen metabolism (SULT1E1, UGT1A1, CYP1A2, ERBB2), fatty acid synthesis (ELOVL2, ELOVL4, ELOVL7), and amino acid transporter (SLC38A1, SLC38A3). High expression of these genes corresponds to poor survival of EC patients. On the other hand, genes involved in glycerophospholipid metabolism (LPCAT2, PLA2G2C, PLA2G2D, PLA2G10, GPCPD1, and PLPP2) are downregulated in cluster-2 samples, and their low expression is correlated with poor survival outcomes. We also identified GCK, PSAT1, GLDC, and UGT3A1 as prognostic markers for EC. They are DEGs in both normal vs. tumor and cluster-1 vs. cluster-2 comparisons.

### Genomic alterations of metabolic clusters

Next, we utilized somatic mutation and CNV datasets to characterize each cluster at the genome level. The somatic mutation dataset for cluster-1 and cluster-2 consists of 300 samples (out of 309) and 225 samples (out of 233), respectively. **Table S2** summarizes the mutation profile of cluster-1 and cluster-2 samples. The mean Tumor Mutation Burden (TMB) per Mb of cluster-2 (32.38) is higher compared to cluster-1 (25.78) samples. The five most frequently mutated genes in cluster-1 samples are PTEN (90%), AR1D1A (56%), PIK3CA (55%), TTN (40%), and PIK3R1 (39%) (**Figure 5a**). On the other hand, TP53 (70%), PIK3CA (44%), TTN (39%), PTEN (32%), and PPP2R1A (30%) are the top five frequently mutated genes in cluster-2 samples (**Figure 5b**). TTN is frequently mutated in both clusters.

**Figure 5:**
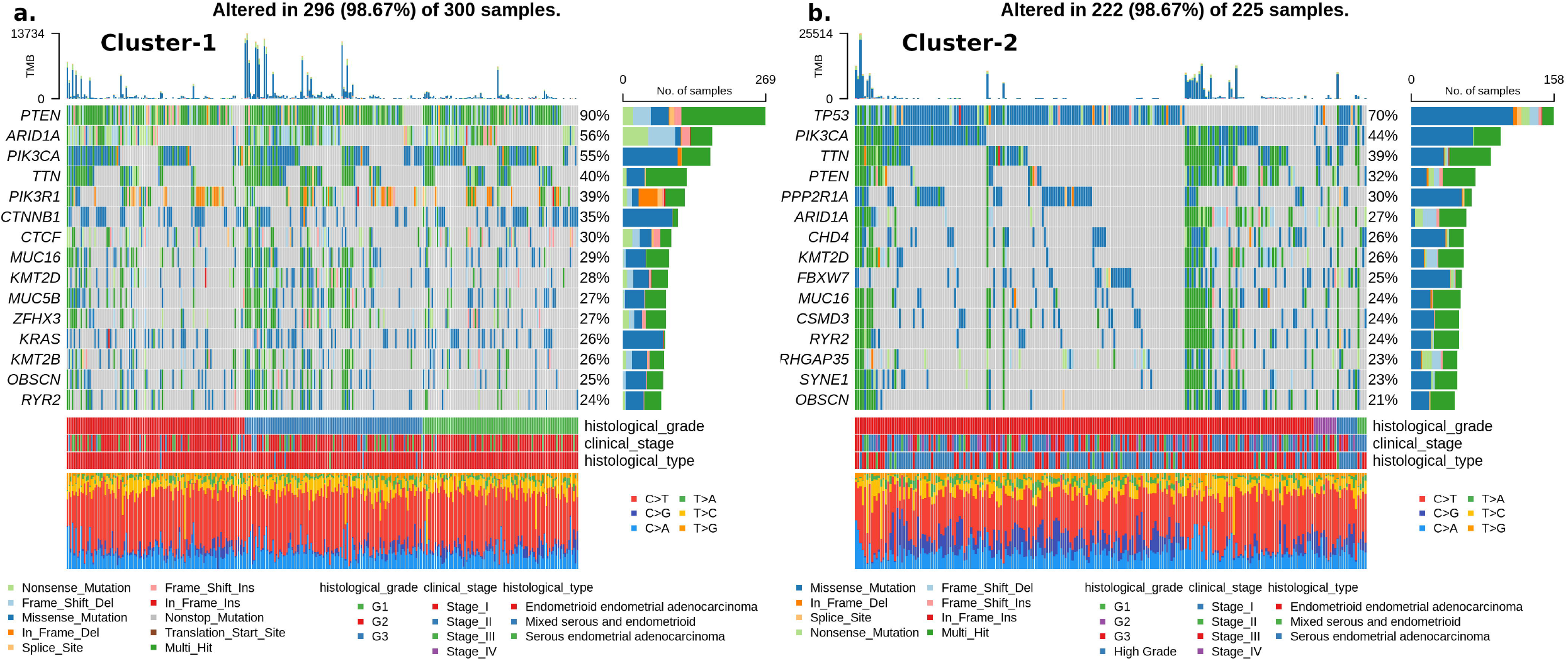
Oncoplot showing frequently mutated genes in (a) cluster-1 and (b) cluster-2 samples. Clinical subtypes and stages of samples are given along with various mutation type.

We compared the mutation profiles of clusters and identified 72 differentially mutated genes (p-value < 0.01) between the clusters. Out of 72, 16 genes are enriched in cluster-1, while 56 are differentially mutated in cluster-2 samples. Interestingly, we found four metabolic genes (UBIAD1, PDK1, GOT2, CYP1A1) to be enriched in cluster-2 samples. UBIAD1 plays a crucial role in cholesterol and phospholipid metabolism. It has tumor suppressor function, and its loss leads to tumor progression with elevated cholesterol levels [33]. GOT2 encodes a mitochondrial enzyme that maintains aspartate-malate shuttle and redox balance and helps cancer cells to proliferate by raising the intracellular aspartate level [34]. CYP1A1, a key enzyme of estrogen metabolism, is also associated with breast cancer proliferation and survival [35]. The forest plot shows the differentially mutated genes, including PTEN, TP53, PPP2R1A, AR1D1A, CTNNB1, PIK3R1, KRAS, CTCF, GOLGA8, and CSNK1A1 (**Figure 6a**).

**Figure 6:**
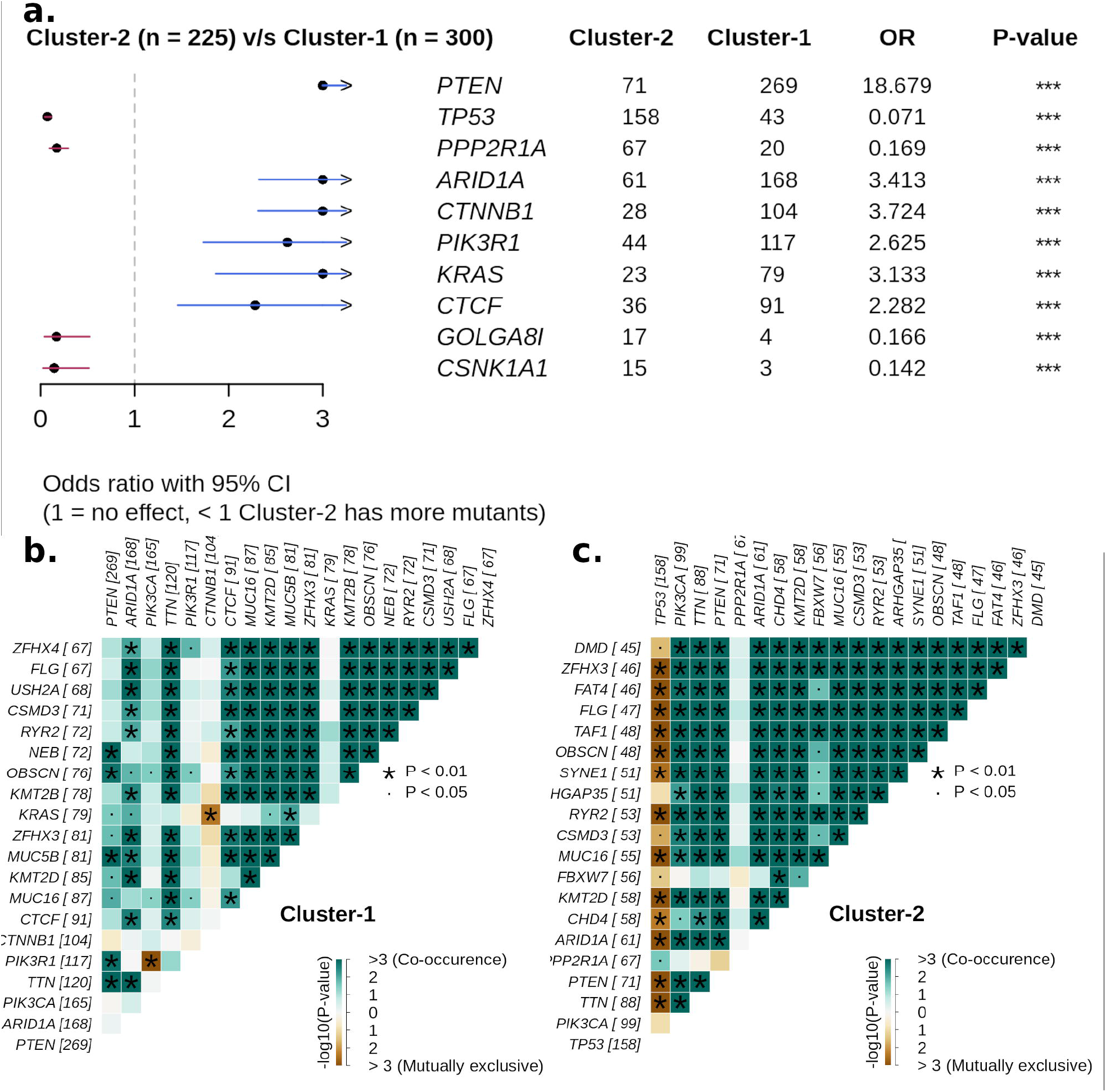
(a) Forest plot showing differentially mutated genes between cluster-1 and cluster-2 samples. (b) Co-occurrence analysis between frequently mutated genes of cluster-1 samples. (c) Co-occurrence analysis between frequently mutated genes of cluster-2 samples.

Cluster-2 includes samples with TP53 mutation, while cluster-1 has PTEN mutations. To further investigate, we analyzed the co-occurrence/mutual exclusive relationship of the 20 frequently mutated genes in both clusters. **Figure 6c** shows that TP53 and PTEN exhibit mutually exclusive behavior (p-value < 0.01). TP53 is also mutually exclusive with other frequently mutated genes in cluster-2 samples except for PPP2R1A. It has also been reported that PPP2R1A mutation is associated with poor survival in breast cancer, lung cancer, and melanoma [36]. The co-occurrence of PTEN along with OBSCN, ZFHX3, KMT2D, MUC16, and TTN is observed in cluster-1 and cluster-2 samples (**Figure 6b-c**). Further, we performed a survival analysis of the frequently mutated genes to reveal their impact on the survival outcomes of EC patients. Among them, six genes have a significant (p-value < 0.05) association with survival (**Table S3**). Patients with PTEN mutation have better survival than those without PTEN mutation (**Figure 7**). Similarly, other genes AR1D1A, MUC16, PIK3CA, and CTNNB1, are related to better survival (Hazard ratio < 1) of EC patients (**Figure 7**). In contrast, patients with TP53 are significantly associated with poor survival.

**Figure 7:**
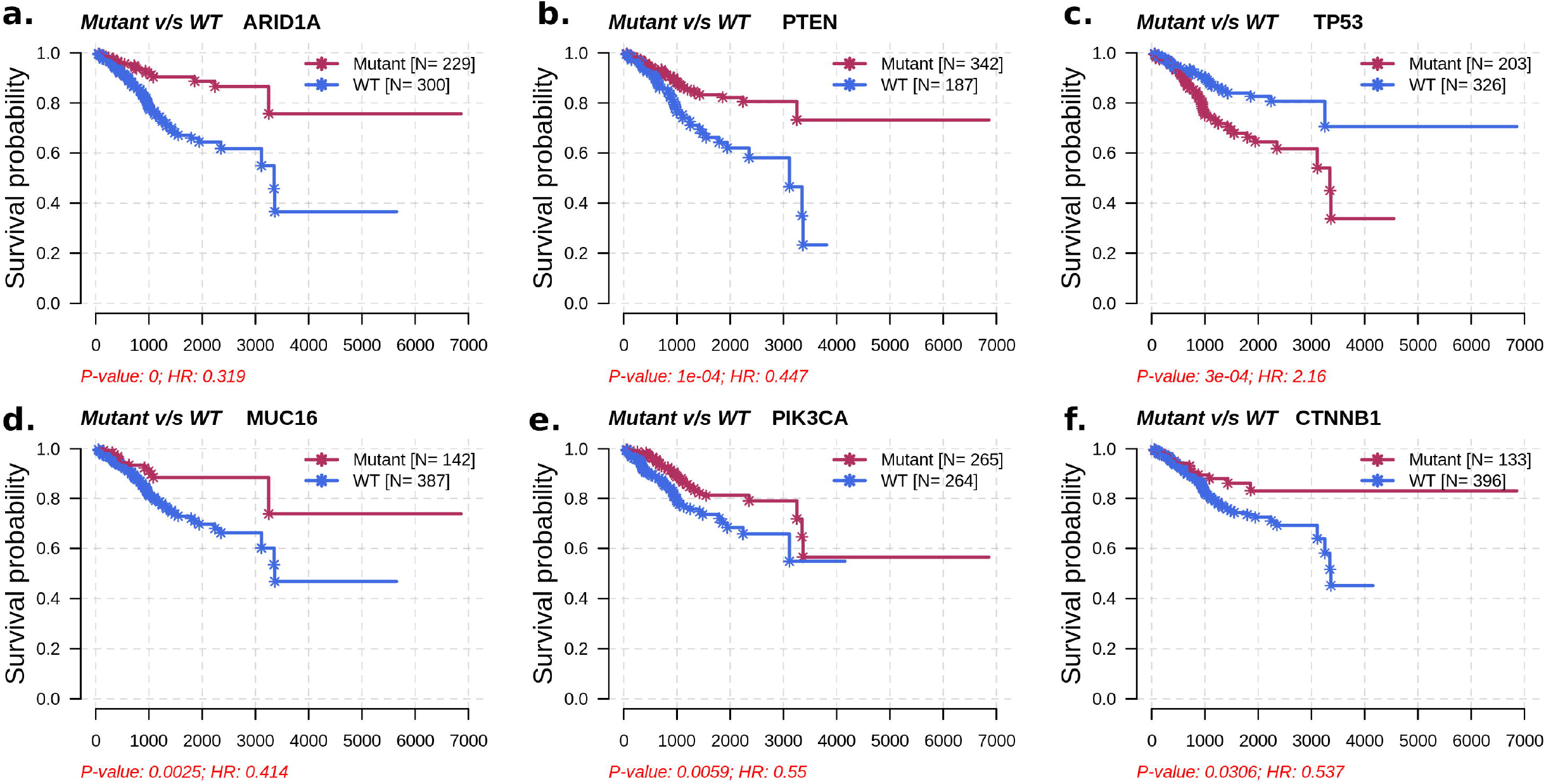
Survival analysis of frequently mutated genes in EC based on their mutation status. HR represents Hazard’s Ratio.

The CNV datasets consist of 534 samples, with 303 samples in cluster-1 and 231 in cluster-2. We employed GISTIC 2.0 to examine the CNVs in each cluster and identified 27 (10 amplified and 17 deleted) and 108 (53 amplified and 55 deleted) significant CNVs in cluster-1 and cluster-2 samples, respectively. These CNVs of cluster-1 and cluster-2 mapped to 23 and 103 chromosome loci, respectively. There are 19 common chromosome loci in both cluster-1 and cluster-2 samples. We observed that cluster-1 samples have very few CNVs (amplifications and deletions) compared to cluster-2 samples (**Figure 8a**). The genome plot shows the chromosome locus of the top 10 significant CNVs (sorted based on q-value) of cluster-1 and cluster-2 samples (**Figures 8b-c**). Out of these, we found 4 chromosome locations to be only altered in cluster-2 samples: 19p13.3, 17q12, 5q12.1, and 4q34.3. Two CNVs corresponding to chromosome locus 19p13.3 are deleted in more than 50% of cluster-2 samples. Frequent deletion of 19p13 has been observed in ovarian cancer and metastatic melanoma [37,38].

**Figure 8:**
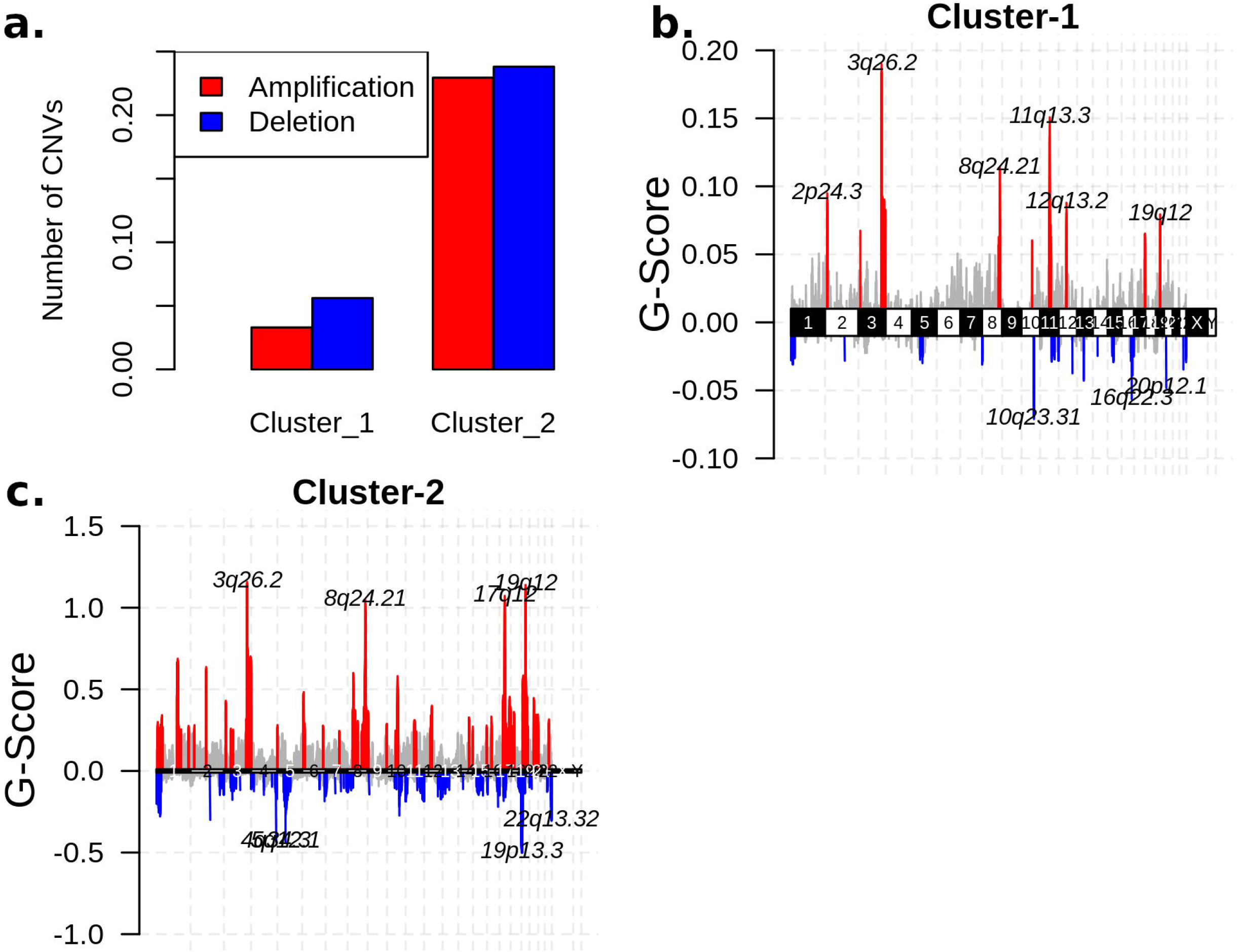
(a) The average number of significant CNVs in cluster-1 and cluster-2 samples. The genome plot showing the G-score of significant CNVs of (b) cluster-1 and (c) cluster-2 samples for all chromosomes. The red and blue colors represent the amplifications and deletions, respectively. The grey color represents the non-significant CNVs.

Further, the significant CNVs of cluster-1 and cluster-2 samples map to 1567 and 4264 genes, respectively. Interestingly, we found 11 metabolic gene alterations that also show consistent changes at the transcriptomic level in cluster-2 samples. These include eight amplified genes (PNMT, ERBB2, ZDHHC19, OXCT2, CARNS1, SLC38A3, SLC6A19, SLC6A3) and three deleted genes (DPEP1, GCNT3, FTMT) that are upregulated and downregulated in cluster-2 samples, respectively. PNMT, a key enzyme of catecholamine (CAT) synthesis, catalyzes the synthesis of adrenaline from noradrenaline by transferring the methyl group from SAM. We also found PNMT as a prognostic marker for EC. PNMT, along with the ERBB2 gene, is located on chromosome 17q12, which is amplified in 35% of cluster-2 samples. ERBB2 controls the activation of estrogen receptors (ESR1 and ESR2) and regulates estrogen metabolism. ZDHHC19 is amplified in more than 60% of cluster-2 samples. It is an oncogene involved in leukotriene metabolism and is present on chromosome locus 3q29 along with PPP1R2. OXCT2 plays an essential function in ketone body catabolism by converting fatty acids to ketone bodies. CARNS1(11q13.2), SLC38A3 (3p21.31), and SLC6A19 (5p15.33) are involved in the synthesis of carnosine, transportation of glutamine, and neutral amino acid, respectively. The high expression of dopamine transporter SLC6A3 (5p15.33) has been observed in multiple cancers [39,40]. DPEP1 (16q24.3) is deleted in more than 60% of cluster-2 patients, and it catalyzes the conversion of leukotriene D4 to leukotriene E4. FTMT (5q12.3) regulates the iron levels in mitochondria and cytosol. The high expression of this gene affects cellular iron homeostasis and inhibits the proliferation of cancer cells [41]. GCNT3 (15q14) is also deleted in cluster-2 samples (>50%). It plays an essential role in mucin biosynthesis and is a prognostic marker in EC.

### Validation of Metabolic Subtypes

We used independent GEO microarray data to validate our results. Out of 1000 MAD genes used for clustering, 932 metabolic genes that overlap with the GEO dataset were considered for NMF clustering with identical parameters. We obtained two metabolic subtypes with cophenetic and silhouette coefficients of 0.98 and 0.91, respectively, which displayed consistency with the TCGA cohort (**Figure S2a**). Metabolic subtypes exhibit a significant correlation (p-value < 0.05) with histological types and grade. Further, the DEGs identified between subtypes overlap with candidates from the TCGA cohort and show similar expression patterns (**Figure S2b**). The expression pattern of some of the candidate genes is shown in **Figure 9**.

**Figure 9:**
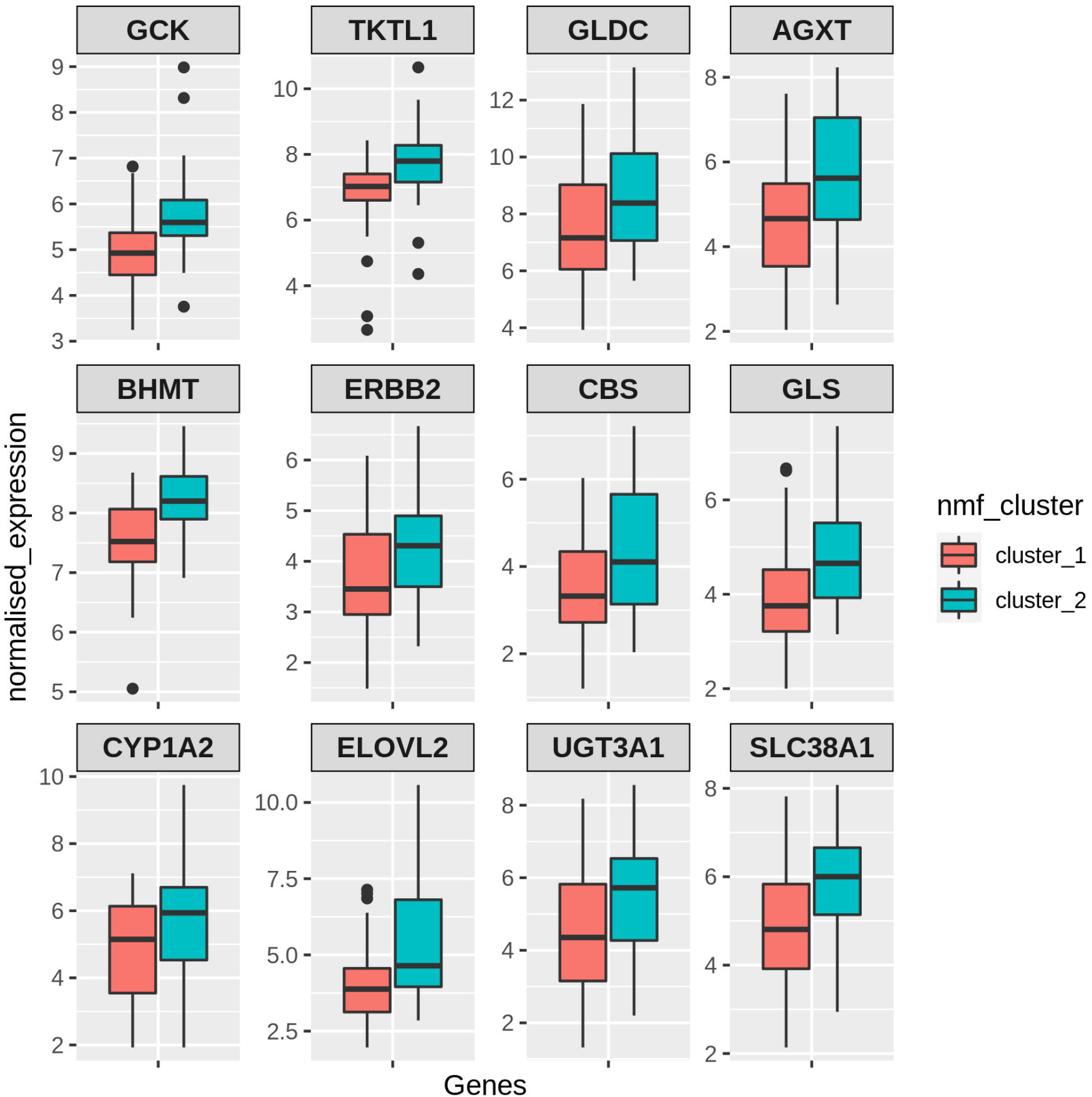
Boxplot showing the expression pattern of candidate genes between two metabolic clusters identified in the GEO validation dataset.

## Discussion

Previous studies demonstrated that cancer tissues exploit different metabolic pathways to produce energy and biomolecules along with maintaining redox homeostasis [12,13]. The utilization of various mechanisms depends on intrinsic and extrinsic factors which contribute to heterogeneity in the metabolic profiles of cancer patients. Earlier studies on EC analyzed the expression of metabolic genes between normal and tumor samples based on differential gene expression analysis [42,43]. In this study, we applied an unsupervised technique to stratify EC samples into metabolic subtypes based on a genome-scale metabolic model (HMR2.0) and characterized each subtype with distinct survival outcomes, clinical features, metabolic pathways, and genomic alterations. We also compared the metabolic dysregulation of subtypes with changes between normal and tumor conditions. In addition, we leveraged the transcriptomic data to extract reporter metabolites for distinguishing subtypes and normal and tumor conditions.

The consensus clustering of metabolic genes of HMR2.0 based on transcriptomic data of EC revealed two robust metabolic subtypes that showed significant correlation to clinical characteristics, including survival, tumor stages, grades, and histological types (**Figures 1** and **2**). Our analysis of differences between subtypes showed the co-activation of the PPP along with one-carbon metabolism in cluster-2 samples with poor survival characteristics (**Figures 3** and **4**). This intersection of metabolic pathways may facilitate the balance between the production of nucleotides and antioxidant species, which support a high proliferation rate and provide defense against oxidative stress [44]. Accordingly, we also observed glycolytic genes to be upregulated in EC, which is consistent with previous studies [17,42]. The candidate genes of PPP include TKTL1, which is upregulated in cluster-2 samples. The transketolase enzyme is part of the non-oxidative branch of PPP that plays a crucial role in the ribose synthesis required for nucleotides. TKTL1 protein is also shown to be overexpressed in EC [45]. It is related to poor survival in different cancers, and inhibition of TKTL1 leads to a significant reduction in the proliferation of many cancers [46]. GCK, a prognostic candidate gene of the hexokinase family, is upregulated in Cluster-2 samples, and its high expression is associated with poor survival.

The serine synthesis pathway is upregulated in cluster-2 samples (**Figure 4**). We observed metabolites of one-carbon metabolism as reporter metabolites. Overexpression of genes involved in the serine biosynthesis pathway is also related to poor survival in breast cancer [44]. Serine and glycine provide one-carbon units for nucleotide synthesis through the folate cycle and contribute toward mitochondrial NADPH production. We found folate receptor FOLR3 to be upregulated in cluster-2 samples, and its overexpression has been reported in ovarian cancer (**Figure 3a**) [47]. One-carbon metabolism also contributes to the production of SAM, which is a reporter metabolite in EC samples. SAM is used in the methylation of biomolecules through the methionine cycle. In cluster-2 samples, we observed choline and betaine as reporter metabolites used in the production of methionine from homocysteine (**Figure 4**). These are unreported changes in EC, and the upregulation of BHMT is associated with poor survival. Serine can also be used in the production of cysteine (through the transsulfuration pathway), which is utilized in the synthesis of glutathione. CBS, involved in the first step of the transsulfuration pathway, is upregulated in cluster-2 samples and is associated with poor survival. The elevated expression of CBS is responsible for inducing tumor growth in other gynaecological cancers, including ovarian and breast cancer [48]. The changes in one-carbon metabolism may mirror the DNA methylation frequency observed in high-grade tumors [49].

The prognostic candidates include genes of estrogen metabolism. EC is characterized by elevated estrogen exposure. Upregulation of SULT1E1 can inactivate the estrogens in cluster-2 samples and contribute to the difference in estrogen levels of both subtypes. In cluster-2 samples, we observed high expression of ERBB2, which is correlated with poor survival of EC patients [50]. GLS, an essential gene involved in the hydrolysis of glutamine, is also upregulated in cluster-2 samples. It has been shown that GLS in EC is regulated by estrogen [51].

Although only metabolic gene expression data was used for clustering, subtypes identified are described by distinct mutation and CNV patterns capturing the relationship between genomic alterations and metabolic gene expression (**Figures 5** and **6**). Cluster-1 samples are associated with PTEN mutation and low CNVs, whereas cluster-2 samples are associated with TP53 mutation and high CNVs. Both PTEN and TP53 are tumor suppressor genes, and their suppressor functions are associated with cytoplasm and nucleus, respectively [52]. PTEN and TP53 also function as regulators of various metabolic processes to maintain cellular homeostasis [53,54]. We observed a mutually exclusive pattern between PTEN and TP53 mutation in cluster-2 samples. Kurose et. al., (2002) showed that PTEN and TP53 mutations are mutually exclusive in breast cancer [55]. We also observed that PTEN and TP53 mutations have opposite survival outcomes in EC (**Figure 7**). Somatic mutation in PTEN is associated with better survival, while TP53 mutation is related to poor survival. This finding is consistent with the observation made by Risinger et al., (1998) [56]. ARID1A mutation exists in the preneoplastic stages, and its loss may not be enough to develop EC [57]. In addition, PNMT and ERBB2 are upregulated, amplified, and present in the same chromosome locus, 17q12, in cluster-2 samples (**Figure 8c**). We identified PNMT and ERBB2 as prognostic genes for EC. Gene amplification and coexpression of PNMT and ERBB2 have been observed in breast cancer patients [58]. Further, 17q12 amplification is related to high-grade breast cancer and has been associated with poor outcomes [59].

In the TCGA study, Levine et al., (2013) identified four clusters using CNV data of 363 EC patients [10]. The TCGA cluster-1 samples consist of a few genomic alterations, while cluster-2 and cluster-3 samples are distinguished by 1q amplification. Cluster-4 comprises many amplified chromosome locations, including 8q11.23 (SOX17), 19q12 (CCNE1), and 4p16.3 (FGFR3) and deletion of LRP1B. In our study, we observed very few significant CNVs in cluster-1 samples and amplification of 1q, 8q11.23 (SOX17), 19q12 (CCNE1), and 4p16.3 (FGFR3) in cluster-2 samples. These observations indicate that metabolic cluster-1 has the characteristics of TCGA cluster-1 and cluster-2 samples based on CNV data, while cluster-2 has the characteristics of TCGA cluster-3 and cluster-4 samples. Additionally, cluster-1 samples exhibit characteristic features of Type 1 EC, while cluster-2 samples have the characteristics of Type 2 EC based on the association of clusters with survival outcomes, mutation status, and distinct CNVs.

## Conclusions

Our study revealed the metabolic heterogeneity within EC and identified subtypes with distinct transcriptomic, genomic, and clinical features. Although there are fewer mutations, amplifications, and deletions in metabolic genes, the metabolic changes observed at the transcriptome level are associated with the overall genomic changes observed in EC. A comprehensive elucidation of distinct dysregulated metabolic pathways, which correlate with clinical outcomes, will assist in diagnosing and treating EC.

## Abbreviations

TCGA: The Cancer Genome Atlas
GEO: Gene Expression Omnibus
HMR2.0: Human genome-scale metabolic model version 2.0
MAD: Median Absolute Deviation
NMF: Non-negative Matrix Factorization
PCA: Principal Component Analysis
DEG: Differentially Expressed Gene
FC: Fold Change
TMB: Tumor Mutation Burden
CNV: Copy Number Variation
GISTIC: Genomic Identification of Significant Targets in Cancer
PPP: Pentose Phosphate Pathway
TCA: Tricarboxylic Acid

## Declarations

### Ethics approval and consent to participate

Not applicable

### Consent for publication

Not applicable

### Availability of data and materials

The datasets analysed in the current study are available from public repositories: https://portal.gdc.cancer.gov/ and https://www.ncbi.nlm.nih.gov/geo/

### Authors’ contributions

Conceptualization: PKV; Methodology: AS; Formal analysis and investigation: AS; Writing - original draft preparation: AS; Writing - review and editing: AS, PKV; Funding acquisition: P KV; Supervision: PKV

## Acknowledgements

Not applicable

## Figure Legends

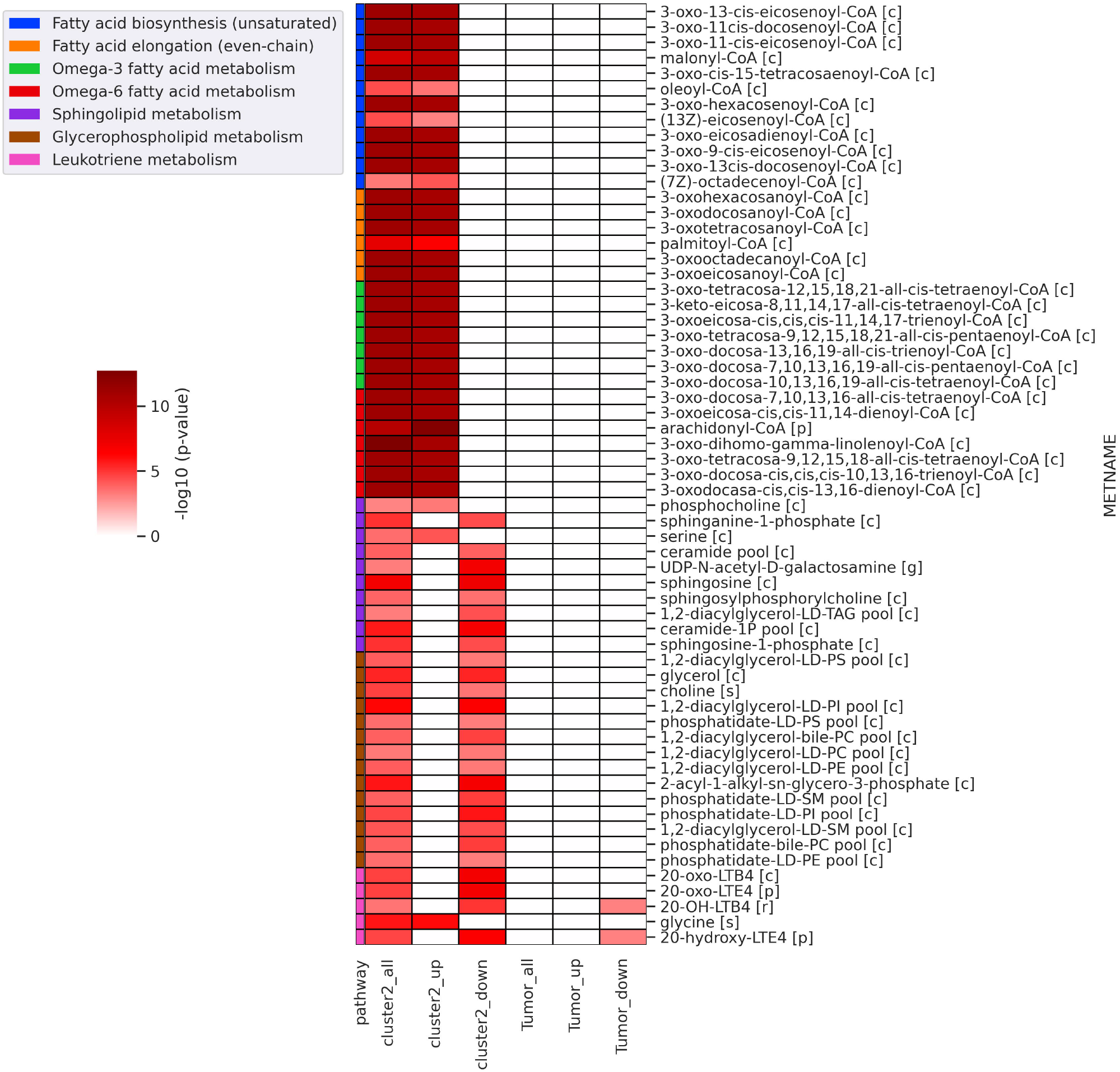

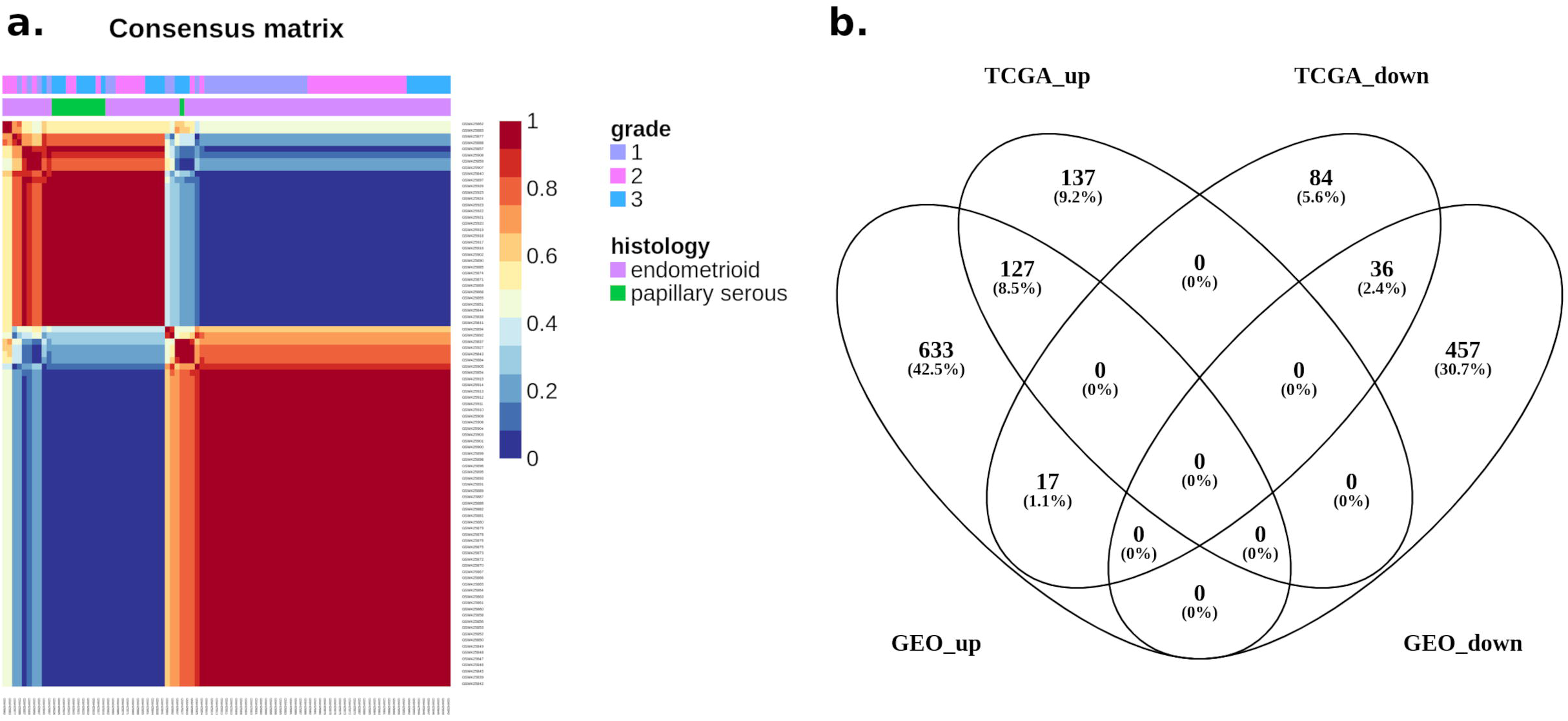

