## Supplementary Material for "Identification and characterization of metabolic subtypes of endometrial cancer using systems-level approach"

### Supplementary Tables

**Table S1**: Fisher exact test and Cramer's V to determine and quantify the association of clusters with clinical variables.

| **Clinical Variables** | **p-value** | **Cramer's V** |
|---|---|---|
| Clinical Stage | 4.117 e-10 | 0.361 |
| Histological Types | 2.2 e-16 | 0.614 |
| Histological Grade | 2.2 e-16 | 0.585 |
| Age (>=50) | 0.0269 | 0.098 |

**Table S2**: Summary of average mutation types in cluster-1 and cluster-2 samples

| **Mutation types** | **Cluster-1** | **Cluster-2** |
|---|---|---|
| Frame_Shift_Del | 38.42 | 26.018 |
| Frame_Shift_Ins | 14.997 | 10.72 |
| In_Frame_Del | 3.287 | 2.396 |
| In_Frame_Ins | 0.207 | 0.191 |
| Missense_Mutation | 769.017 | 1006.698 |
| Nonsense_Mutation | 77.157 | 86.222 |
| Nonstop_Mutation | 0.767 | 0.844 |
| Splice_Site | 18.52 | 25.431 |
| Translation_Start_Site | 0.7 | 0.907 |
| TMB | 923.07 | 1159.427 |
| TMB per mb | 25.78 | 32.38 |

**Table 3**: Survival analysis based on mutation status of frequently mutated genes in EC. Genes with p-value < 0.05 have a significant association with survival.

| Genes | p_value | Hazard's Ratio (HR) | Wild Type (WT) | Mutant |
|---|---|---|---|---|
| ARID1A | 3.77E-06 | 0.319 | 300 | 229 |
| PTEN | 0.000131 | 0.447 | 187 | 342 |
| TP53 | 0.000303 | 2.16 | 326 | 203 |
| MUC16 | 0.0025 | 0.414 | 387 | 142 |
| PIK3CA | 0.00591 | 0.55 | 264 | 265 |
| CTNNB1 | 0.0306 | 0.537 | 396 | 133 |
| ZFHX3 | 0.0636 | 0.6 | 402 | 127 |
| KMT2D | 0.0672 | 0.617 | 385 | 144 |
| TTN | 0.116 | 0.697 | 319 | 210 |
| PIK3R1 | 0.975 | 1.01 | 367 | 162 |

#### Supplementary Figures

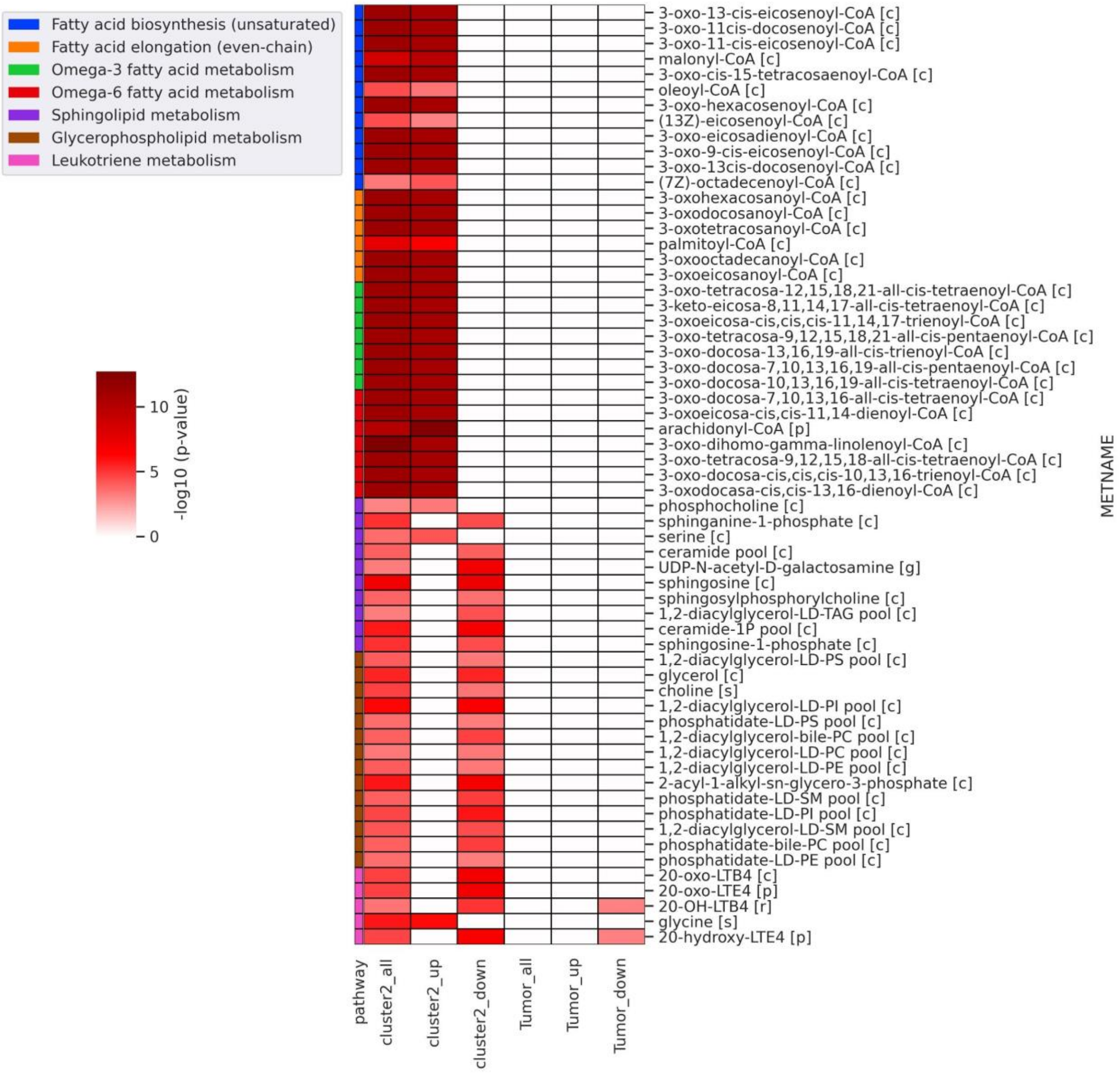


**Figure S1**: Heatmap of significant reporter metabolites in lipid metabolism obtained for different conditions (cluster2_all: all DEGs of cluster-1 vs. cluster-2 condition, cluster2_up: upregulated genes in cluster-1 vs. cluster-2 condition, cluster2_down: downregulated genes in cluster-1 vs. cluster-2 condition, tumor_all - all DEGs of normal vs. tumor conditions, tumor_up: upregulated genes in normal vs. tumor conditions, tumor_down: downregulated genes in cluster-1 vs. cluster-2 conditions). The metabolite name (METNAME) also includes the compartment information, [c] - cytosol, [m] - mitochondria, [n] - nucleus, [p] - peroxisome, [g] - Golgi apparatus, [s] - extracellular, [r] - endoplasmic reticulum.

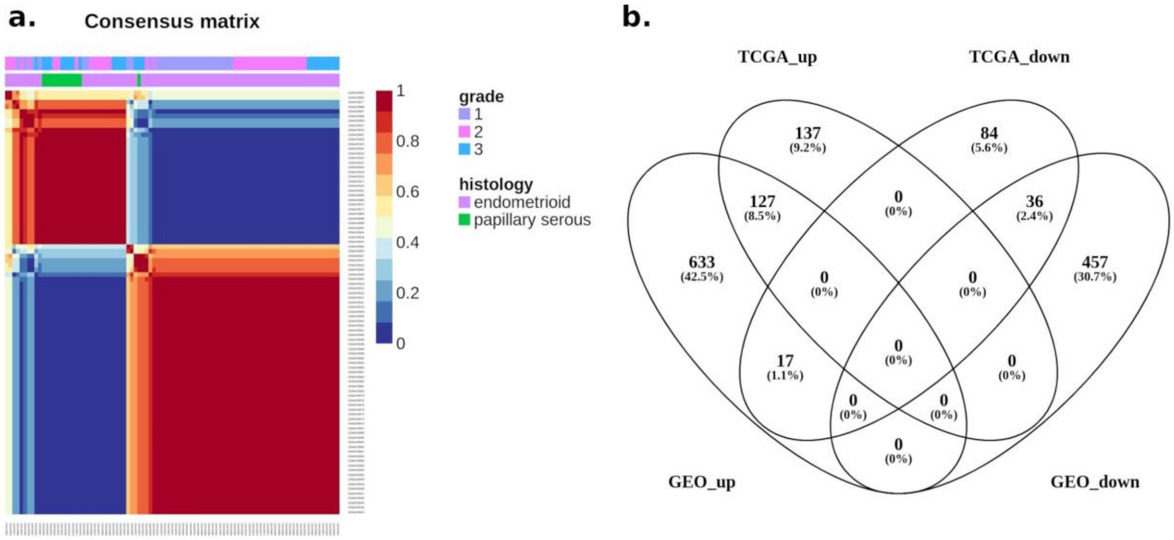


**Figure S2**: Validation of metabolic subtypes in an independent GEO dataset. (a) The consensus plot showing that tumor samples are clustered into two groups. (b) Comparison of DEGs in TCGA and GEO datasets of EC (TCGA_up: upregulated genes between clusters in TCGA cohort, TCGA_down: downregulated genes between clusters in TCGA cohort, GEO_up: upregulated genes between clusters in GEO dataset, GEO_down: downregulated genes between clusters in GEO dataset).
